# A deep learning framework for nucleus segmentation using image style transfer

**DOI:** 10.1101/580605

**Authors:** Reka Hollandi, Abel Szkalisity, Timea Toth, Ervin Tasnadi, Csaba Molnar, Botond Mathe, Istvan Grexa, Jozsef Molnar, Arpad Balind, Mate Gorbe, Maria Kovacs, Ede Migh, Allen Goodman, Tamas Balassa, Krisztian Koos, Wenyu Wang, Norbert Bara, Ferenc Kovacs, Lassi Paavolainen, Tivadar Danka, Andras Kriston, Anne E. Carpenter, Kevin Smith, Peter Horvath

## Abstract

Single cell segmentation is typically one of the first and most crucial tasks of image-based cellular analysis. We present a deep learning approach aiming towards a truly general method for localizing nuclei across a diverse range of assays and light microscopy modalities. We outperform the 739 methods submitted to the 2018 Data Science Bowl on images representing a variety of realistic conditions, some of which were not represented in the training data. The key to our approach is to adapt our model to unseen and unlabeled data using image style transfer to generate augmented training samples. This allows the model to recognize nuclei in new and different experiments without requiring expert annotations.

---

Identifying nuclei is the starting point for many microscopy-based cellular analyses. Accurate localization of the nucleus is the basis of a variety of quantitative measurements, but is also a first step for identifying individual cell borders, which enables a multitude of further analyses. Until recently, the dominant approaches for this task have been based on classic image processing algorithms (e.g. CellProfiler^1^) which were sometimes guided by shape and spatial priors ^2^. A drawback of these methods is the need for expert knowledge to properly adjust the parameters, which typically must be re-tuned when experimental conditions change.

Recently, deep learning has revolutionized an assortment of tasks in image analysis, from image classification ^3^ to face recognition ^4^, and scene segmentation ^5^. It is also responsible for breakthroughs in diagnosing retinal images ^6^, classifying skin lesions with superhuman performance ^7^, as well as incredible advances in 3D fluorescence image analysis ^8^. However, aside from initial works from Caicedo *et al.*^9^ and Van Valen *et al.^10^*, deep learning has yet to significantly advance nucleus segmentation performance.

The 2018 Data Science Bowl (DSB) organized by Kaggle, Booz Allen Hamilton and the Broad Institute, challenged participants to push the state-of-the-art in nucleus segmentation. The goal of the challenge was to develop fully automated and robust methods capable of working in a variety of conditions, including differing cell lines, treatments, and types of light microscopy. The challenge attracted thousands of data scientists from around the world. Approaches using deep learning dominated the competition, achieving scores that shattered what was previously possible: the best performing traditional methods we submitted ranked no higher than 1,000 out of 3,891 submissions in Stage 1. The top deep learning-based methods relied on only a handful of different architectures; the factors that participants commonly believed influenced their method’s ranking were related to data: the amount of data, the pre-processing, and methods used to augment the data.

Our approach was unique among the top-25 submissions in applying image style transfer^11^. It aims to overcome one of the greatest challenges of deep learning, the extent of the annotated training set. We augment the training samples by creating realistic-looking artificial sample images with the texture, coloration, and pattern elements from source images not included in the training set (**Figure 1**).

**Figure 1.**
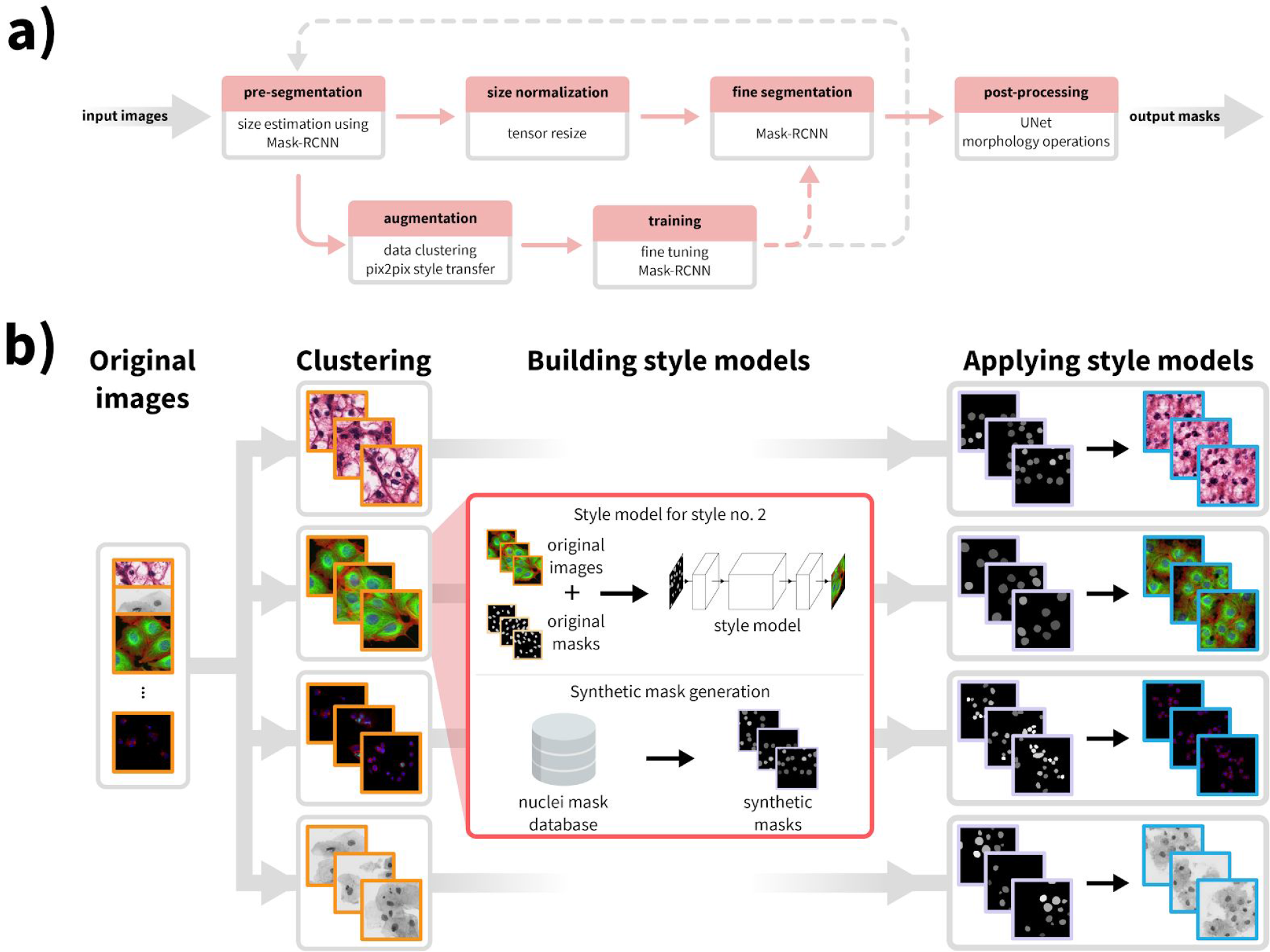
Overview of our approach. a) Upper row of boxes presents the nucleus segmentation and pre-processing; an initial Mask R-CNN network estimates typical nucleus sizes, then images are rescaled such that mean nucleus size is uniform and a Mask R-CNN network trained on images with uniform nucleus size predicts segmentations. A contour refinement step using a U-Net based network with a morphology operation is applied to obtain the final segmentation result. The data augmentation pipeline is depicted in the bottom row, the training set is augmented with an artificially generated set of images similar to the test set, and a pretrained Mask R-CNN method is fine-tuned using the augmented images. Augmentation and training steps may be iteratively repeated as the gray dashed line suggests. Upper row depicts the inference pipeline, bottom row training. Solid lines indicate data flow, dashed lines indicate transfer of a trained model. b) Image style-transfer based data augmentation. To adapt our model to handle image types for which we have no segmentation labels, we synthesize new training data by first clustering images into similar groups, then learning a style transfer model. The style transfer model is provided with simulated nucleus masks which mimic the number, shape, and size of the unseen nuclei, and then synthetic training image/label pairs are generated using the masks and the style transfer models. This data is added to the standard training data provided to Mask R-CNN, and the network learns to segment nuclei in the new domain.

Combining this with a segmentation network based on Mask R-CNN^12^, along with boundary correction using U-Net^13^ (**Supplementary Fig. S4**) and mathematical morphology, our method outperforms all other methods reported on the final Data Science Bowl leaderboard (post-competition)^1^. We also demonstrate that our method outperforms similar baselines on public fluorescent and histology datasets. It does not require parameter tuning or specialized knowledge to use, and can be applied on a wide variety of conditions and imaging modalities.

Our approach (**Figure 1a**, **Figure S1** and **Online Methods)** begins by rescaling the images such that nuclear size is approximately uniform, as we found that the performance of the network is improved if the nuclear size is fixed during training and inference (see **Online Methods** and **Supplementary Figure S3**). To do this, we estimate the typical nucleus size in the provided images with a Mask R-CNN-based network pre-trained on a large set of diverse images with nuclear segmentations, and fine-tuned using the provided training data and label masks. The output of this network is an initial segmentation we use to estimate the typical nucleus size. Alternatively, if the typical cell size is known *a priori*, it can be defined manually and the images rescaled accordingly.

Next, to adapt our model to handle a wide variety of cell types, staining methods, and imaging modalities, even those for which no segmentation annotations are available, we augment the training set with an artificially generated set of representative image/label pairs. This is accomplished using image style transfer. We begin by clustering the images into similar styles based on their appearance (**Figure 1b**). For each cluster of similar image types, a style transfer network^11^ is trained to generate synthetic images of the desired style with nuclei at specified locations. Nucleus annotations are used to train the style transfer network when provided, otherwise we use nucleus masks output from the initial segmentation network. After a style transfer network is trained for each image style, we generate a set of artificial nucleus masks representative of the shape, size and spatial distribution of nuclei belonging to that style. For this, we collected ∼100,000 manually labeled single nucleus masks from the DSB set. A subset of these nuclei is selected that represent the shape distribution of the original morphologies and they are placed such that they follow the spatial distribution of the image style (see **Online Methods**). With trained style networks and representative nucleus masks in hand, we generate synthetic images in the desired style with nuclei in locations defined by the artificial masks. The synthetic image/mask pairs make up the augmented data set; samples are shown in **Figure 1b**. The augmented data is added to the training data. For this experiment, we generated 20 synthetic image/mask pairs for each of the 134 style clusters we identified in the final round data.

Finally, the ultimate Mask R-CNN segmentation model is trained on the combined augmented and rescaled training data. Unseen images are adjusted such that the estimated nuclei size is uniform. To refine the segmentations for high pixel-level accuracy, the edges of each detected nucleus are corrected using a U-Net-based model trained on the same data, followed by some mathematical morphology-based post processing (see **Online Methods**). This step may be skipped if such accuracy is unnecessary for the application, for example if simply counting nuclei.

We evaluated our approach on four different data sets: the DSB Stage 1, DSB Stage 2, a set of fluorescence microscopy images, and a set of histology images from various sources (*DSB1, DSB2, fluo,* and *hist*, respectively). Details of the images are presented in **Supplementary Table S2**. We compare our approach against submissions from other teams on *DSB1* and *DSB2* (nearly 3,000 in Stage 1, and 739 in Stage 2). As benchmarks, we include the results reported in the first and second positions of the leaderboard which was frozen at the close of the competition (https://www.kaggle.com/c/data-science-bowl-2018/leaderboard), a recent deep learning method, unet4nuclei ^9^, a widely used Otsu threshold based method with object splitting ^1^, ilastik ^14^, and a more sophisticated but still classical gradient vector flow (GVF) based method ^15^ (**Fig. 2** and **Supplementary Table S1**). Unet4nuclei is based on a U-Net ^13^ structure, GVF is an active contour-based method that drives the contour to edges using gradient vectors pointing to bright regions, and ilastik is a pixel-based classification software (see **Supplementary Software 2**). Notably, the DSB Stage 2 evaluation is performed on an unknown subset of the provided test images, many of which are outside the domain of the training images, truly challenging the ability of the model to generalize. We provide additional benchmarks and variations of our approach for comparison in **Online Methods** and **Supplementary Figure S2.**

**Figure 2.**
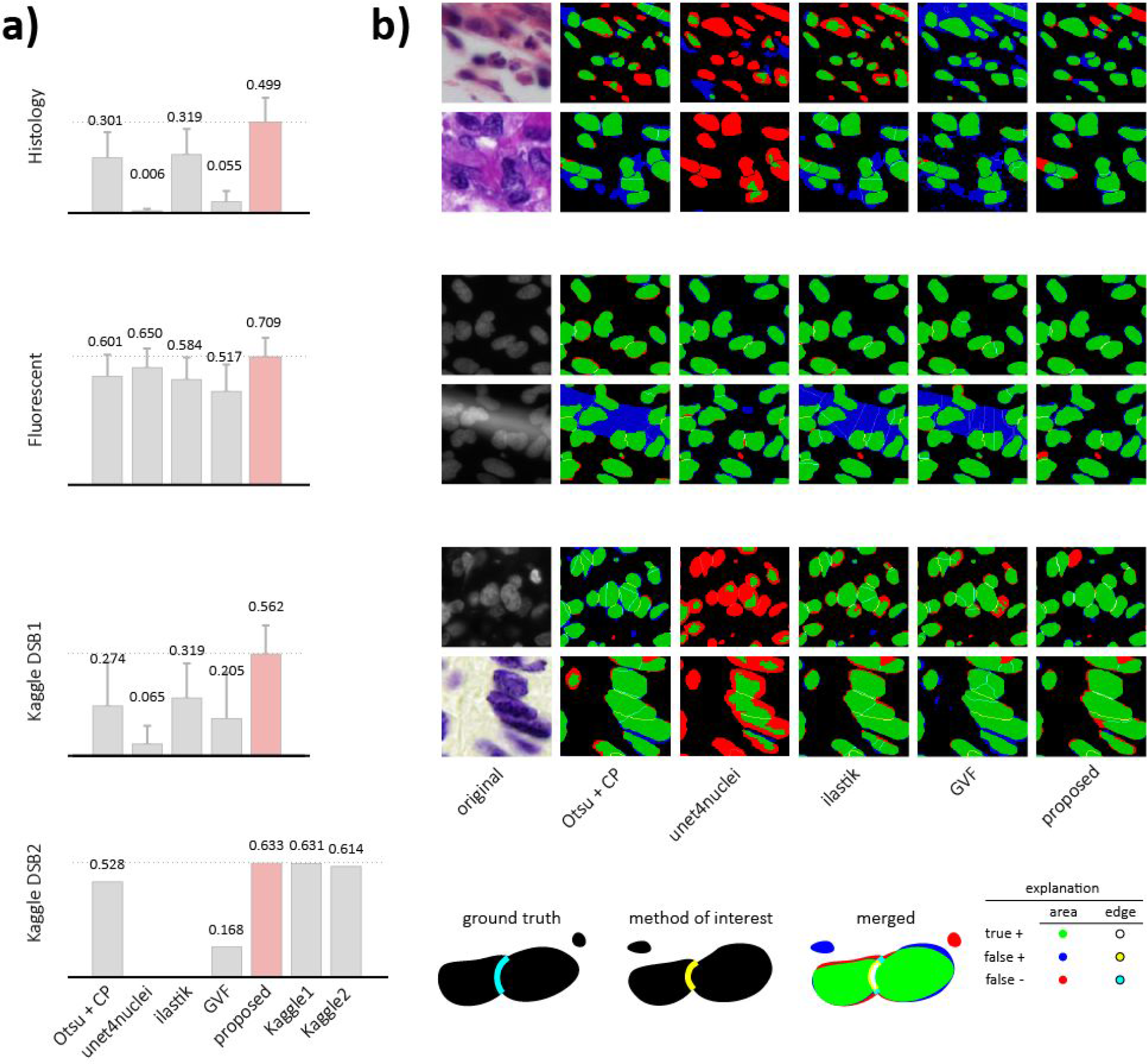
Results. a) DSB scores with error bars for four image sets: Histology, Fluorescent, DSB Stage 1, DSB Stage 2 (see details in **Online Methods**). Highest scores are marked with dashed lines and red color. b) Segmentation results for various methods on sample image crops with difficult cases (two example images of each); rows match those of (a) (note: ground truth is not public for DSB Stage 2). A crop of the original image is provided in the first column, followed by segmentation results predicted by various methods. The color-coding of the results is explained in the legend at the bottom.

The evaluation metric (DSB-score, see definition in **Online Methods**) used in the 2018 Data Science Bowl measures a modified mean average precision^2^ of detected nuclei. For each image, the accuracy of nucleus segmentation is averaged across a set of different intersection-over-union (IoU) thresholds ranging from 0.5 to 0.95 with steps of 0.05. IoU measures segmentation quality as the ratio between the area where the segmented region overlaps a true nucleus and the union of the segmented region and the nucleus.

On the DSB Stage 2 test set, our method scores higher (0.633) than the top ranked (0.631, the highest of 739 teams) and has a simpler architecture with fewer parameters. Results for a traditional Otsu-based baseline segmentation are provided for reference (0.528)^1^]. All *DSB2* results are presented in **Figure 2a**. In addition, our proposed method outperformed all prior published results on *hist*, a diverse set of histology images and on *fluo*, a fluorescent image set (BBBC039 ^9^) (see **Supplementary data** for details). A deeper comparison of our results against six other methods evaluated with additional metrics is provided in **Supplementary Table S1** and **Figure S5**(see details in **Online Methods**).

We proposed a deep learning-based nucleus segmentation approach, using image style transfer to augment our training data with valuable out-of-domain samples. Our segmentation network learned from these artificially-generated image/mask pairs which mimic the patterns of new data types. This approach helped the network adapt to a diverse set of test data outside the domain of the training data, outperforming every other deep learning and classical method tested. Our software is open source and freely available (**Supplementary Software 1** at https://github.com/spreka/biomagdsb). Pre-trained networks for DSB, fluorescent and histology data can be applied to new images via CellProfiler (**Supplementary Software 2**) or through an online interface at www.nucleAIzer.org. The idea of augmenting difficult-to-obtain data using style transfer has enormous potential not only for nucleus detection, but more broadly in bioimage analysis.

## Author contributions

RH., AS., ET., CM., FK., TD., AK., KS., and PH. designed the method. RH., AS., TT., ET., CM., BM., IG., JM., AB., MG., MK., EM., TB., KK., WW., NB., FK., LP., TD., AK., and PH. performed annotation, testing and benchmarking. CM., KK., ET., AK., RH., and PH. designed the online tool. RH., AG., KK., LP., AEC., KS. and PH. wrote the manuscript. AG. and AEC. enabled CellProfiler connection and co-organized the 2018 Data Science Bowl.

## Acknowledgement

We acknowledge support from the LENDULET-BIOMAG Grant (2018-342) and from the European Regional Development Funds (GINOP-2.3.2-15-2016-00006, GINOP-2.3.2-15-2016-00026, GINOP-2.3.2-15-2016-00037), the Academy of Finland grant 310552 to LP. Funding was provided by the National Institute of General Medical Sciences of the National Institutes of Health under MIRA R35 GM122547 to AEC. Funding was provided by the Swedish Research Council (Vetenskapsrädet) under research project grant 2017-040609_3 to KS. We gratefully acknowledge the NVIDIA Corporation for the donation of NVIDIA TITAN Xp GPUs for our research.

## Online Methods

### 1. Overview of the pipeline

As a first step, pre-segmentation of the input images is performed using a pre-trained deep convolutional model (which we refer as *preseg*) to estimate nuclei sizes as well as to create a mask input for image style transfer learning. Simultaneously, we cluster similar images of the input data into groups, and learn styles on these clusters (see Fig.1.b and 2.1.2-2.1.3 for details). As a next step, we extend the training data with artificially created style transferred images for fine-tuning a Mask R-CNN ^12^ pre-trained on our nucleus segmentation dataset. For inference on unseen data, we use the refined Mask R-CNN network incorporating knowledge about estimated cell sizes. The resulting contours are refined with U-Net ^13^ and a morphology step.

**Figure S1.**
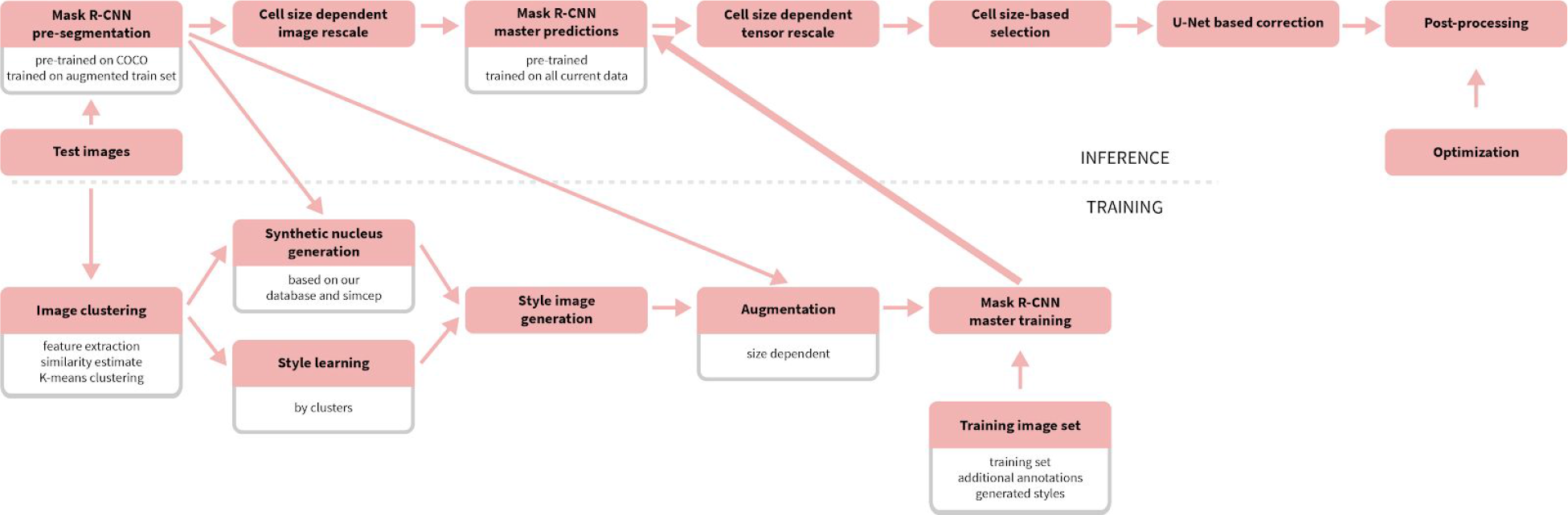
A detailed overview of the proposed pipeline.

### 2. Details of implementation

The proposed method consists of procedures for training and inference, as shown in **Fig. S1**. Inference merely requires unannotated images as its input – provided the pre-trained models are available. Training the network produces a learned segmentation model, and requires a set of annotated training data and a pre-trained segmentation network (pre-segmentation network), as well as any available unannotated images that can be used for data augmentation. The pretrained segmentation network is crucial to both the training and inference procedures, so we discuss it first and then continue with training and prediction steps.

#### 2.1. Training and style transfer data augmentation

##### 2.1.1. Pre-segmentation

The architecture for the segmentation networks is based on the Mask R-CNN architecture. The pretrained segmentation network (pre-segmentation network) is used to make rough estimates about the nucleus size and shape while being robust to changes in imaging modality or magnification. The network is initialized with pretrained weights from the MS-COCO dataset, which contains images and segmentation masks for 91 object types including people, trucks, sheep, dogs, etc. For details about the original COCO competition see http://cocodataset.org or the corresponding publication^16^ (Lin et al. 2014, arXiv). The network was trained using a diverse set of annotated images containing various imaging modalities, cell lines, magnifications, etc. For more information see Section 3.2. The annotations consisted of segmentation masks for the nuclei. Augmentation was used during training including geometric transformations, intensity stretching, cropping, noise, and blur (see **Supplementary Software 1** documentation for details).

The resulting network, which we refer to as *preseg*, already performed reasonably well on unannotated images in the test set (**Fig. S2**), although this was not its purpose. The preseg network is used to: estimate properties of nuclei in new unannotated images (size, shape, and area) in 2.1.2, and to generate rough segmentations on unannotated images for the style transfer data augmentation step described in 2.1.3.

**Figure S2.**
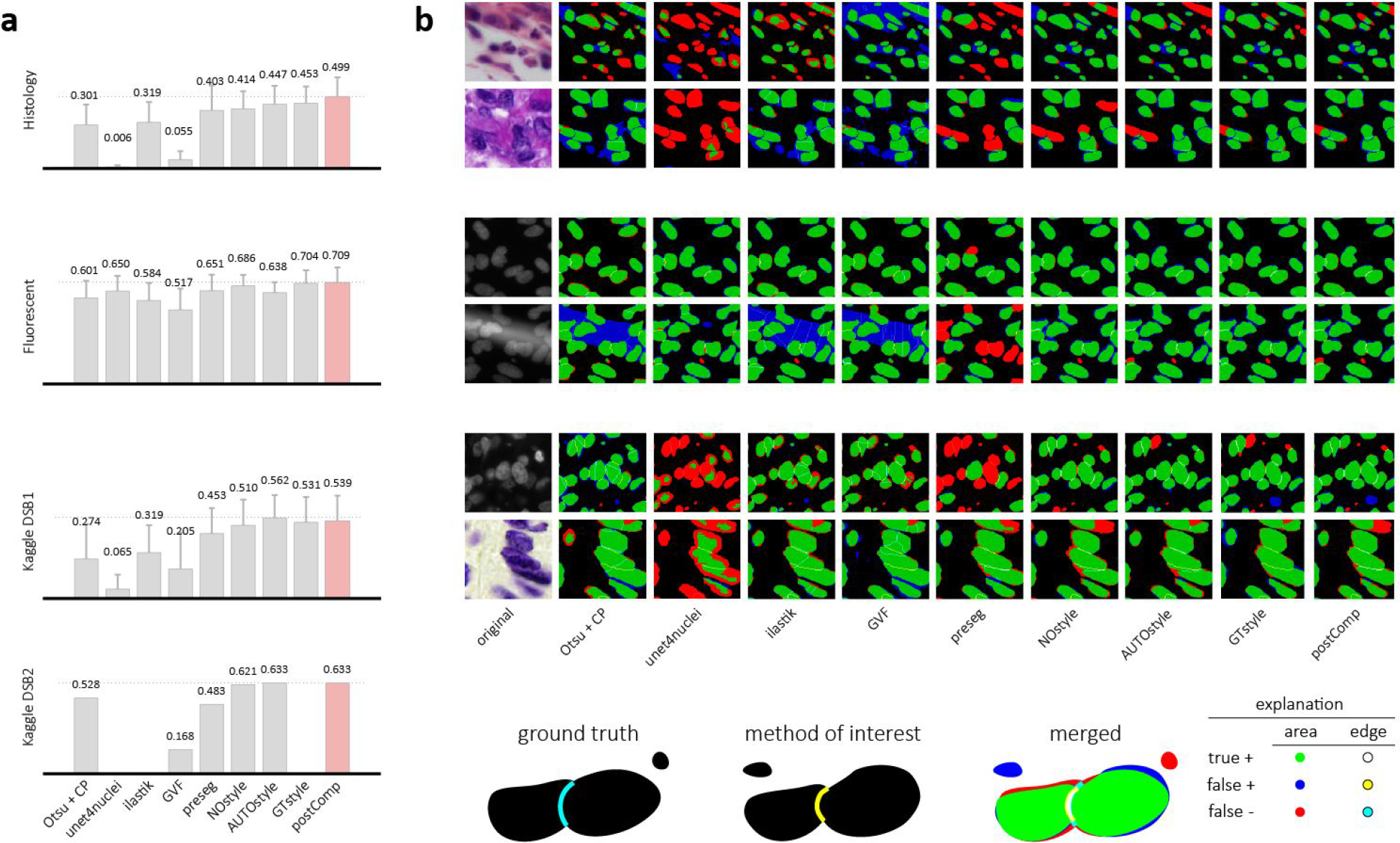
Results from Figure 2 in more detail. a) DSB scores with error bars for the image sets tested. b) Segmentation results for various methods. *preseg*, *NOstyle*, *AUTOstyle*, *GTstyle* and *postComp* refer to variations of our approach. See sections 3.2. and 3.4-3.5. for details.

##### 2.1.2. Clustering for style transfer learning

Images without annotations are automatically clustered to define multiple groups with similar properties: textures, imaging modalities, cell lines, sample type (tissue or culture), etc. These groups are used as data sources to learn style transfer models to generate additional synthetic data that mimics the properties of each cluster of unannotated images.

To perform the clustering, we use a pairwise similarity metric between feature vectors describing each unannotated image. Features were extracted using CellProfiler^1^ modules including intensity and texture and a similarity metric was computed by a shallow fully connected neural network^17^. This similarity network was trained on the DSBtrain1 data set, where images taken with the same condition are given a label of 1 and images from different conditions are given a label of 0. The output of this network on the unannotated data yielded a similarity matrix which we clustered with k-means. The number of clusters, k=134, was chosen to over-segment the groups to avoid accidental mixing of the true underlying groups. Ideally, each obtained cluster of unannotated images represents a “style” or distribution of data which can be augmented with style transfer (e.g. digital slides of H&E stained breast cancer histology samples at 63x magnification, or fluorescent images of Human MCF7 cell nuclei at 40x).

##### 2.1.3 Learning image style transfer models

We use the pix2pix^11^ framework for image style transfer (https://github.com/junyanz/pytorch-CycleGAN-and-pix2pix). The architecture consists of two adversarial networks, a *generator* tasked with synthesizing realistic looking images, and a *discriminator* tasked with identifying real images from synthesized images. This model learns to map one image domain to another through an adversarial loss that encourages the generator to learn to fool the discriminator. The input to the generator in our case is a binary mask containing 1’s at the locations of the desired nuclei, and 0’s elsewhere. The input of the discriminator is an image/mask pair (either a real pair, or a synthetically generated pair). The generator learns to transform the binary mask into the desired style of the real images from the cluster, and the discriminator encourages this by trying to identify real image/mask pairs from fakes. We use the rough segmentations provided by the *preseg* network as masks for the unannotated images in the style cluster during learning. We train a pix2pix style transfer network to synthesize realistic images from masks for each of the style clusters.

##### 2.1.4 Synthesizing new image/mask pairs

Using our set of 134 trained style transfer networks, we synthesized 20 new image/mask pairs for each of the styles in the unannotated data. A crucial step for this task was to generate novel binary masks to provide as input to the style transfer network, which uses the mask to generate a realistic image of the cells with nuclei in the locations defined in the mask. We generated the masks algorithmically as a combination of 1) fetching real nuclei masks from a database, and 2) synthesizing nuclei using software (simcep^18^). Approximately 50% of the nucleus masks were created using each approach. In this manner, we generated 20 masks for each of the 134 style clusters, and then used the style transfer network to generate the corresponding images.

##### 2.1.5. Training the Mask R-CNN segmentation network

The synthetic image/mask pairs generated by the style transfer network were added to the annotated training data to update the Mask R-CNN segmentation network. We used the implementation of Matterport (https://github.com/matterport/Mask_RCNN) and wrote handler scripts in Python to create the appropriate data structures and call functions. Training was performed in 3 steps with decreasing learning rate and targeted different layers of the Mask R-CNN network, as described in the documentation of the aforementioned Matterport repository.

The loss function was as defined in ^12^ (Kaiming He et al): it comprises of classification, localization and segmentation mask losses: L=L_cls_+L_box_+L_mask_ by ROIs, and defines mask loss as follows. Given the *k*-th region does belong to ground truth class *k* it takes the average binary cross-entropy loss which is formulated as

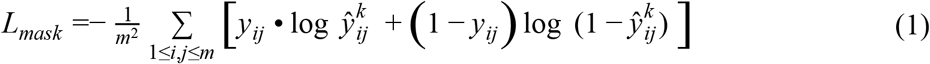

where *y*_*ij*_ is the true label of a cell (*i,j*) from a ROI of *m*x*m* size on the ground truth mask of class *k* and 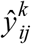 is the predicted class label of the same cell. The formula only includes masks for ground truth class *k* that are associated with the *k*-th class.

##### 2.1.6. Data augmentation and image resizing to uniform nucleus size

The performance of deep learning networks is known to scale with the size of the dataset ^19^. Therefore, we use a number of approaches to augment the training data. The first, as we described above, is to add new synthetic image/mask pairs generated in the style of unseen examples to the existing annotated training data. Each minibatch contained 10-50% synthetic images. We also used standard data augmentation techniques including random cropping, colour channel swapping, intensity modification by histogram stretching or equalization and inversion, rotation to an arbitrary degree and translation as geometric transformations and finally, to better resemble low-quality images, blur and additive noise were used as well. These operations were applied to all the input training data – style transfer results too – with a random probability.

Mask R-CNN is reasonably robust to changes in scale, but superior performance is obtained if the nucleus size is approximately 40 pixels in diameter for the data and parameters we used (**Supplementary Figure S3**). Another preprocessing step was to resize the images by a scaling factor to obtain a training dataset homogeneous both in cell and image size. The scaling factors were computed from the size estimation of the *preseg* nucleus masks such that the resulting mean cell size is set to 40 pixels diameter. Images were then either cropped or padded so that the resulting image was 512 × 512 pixels.

**Figure S3.**
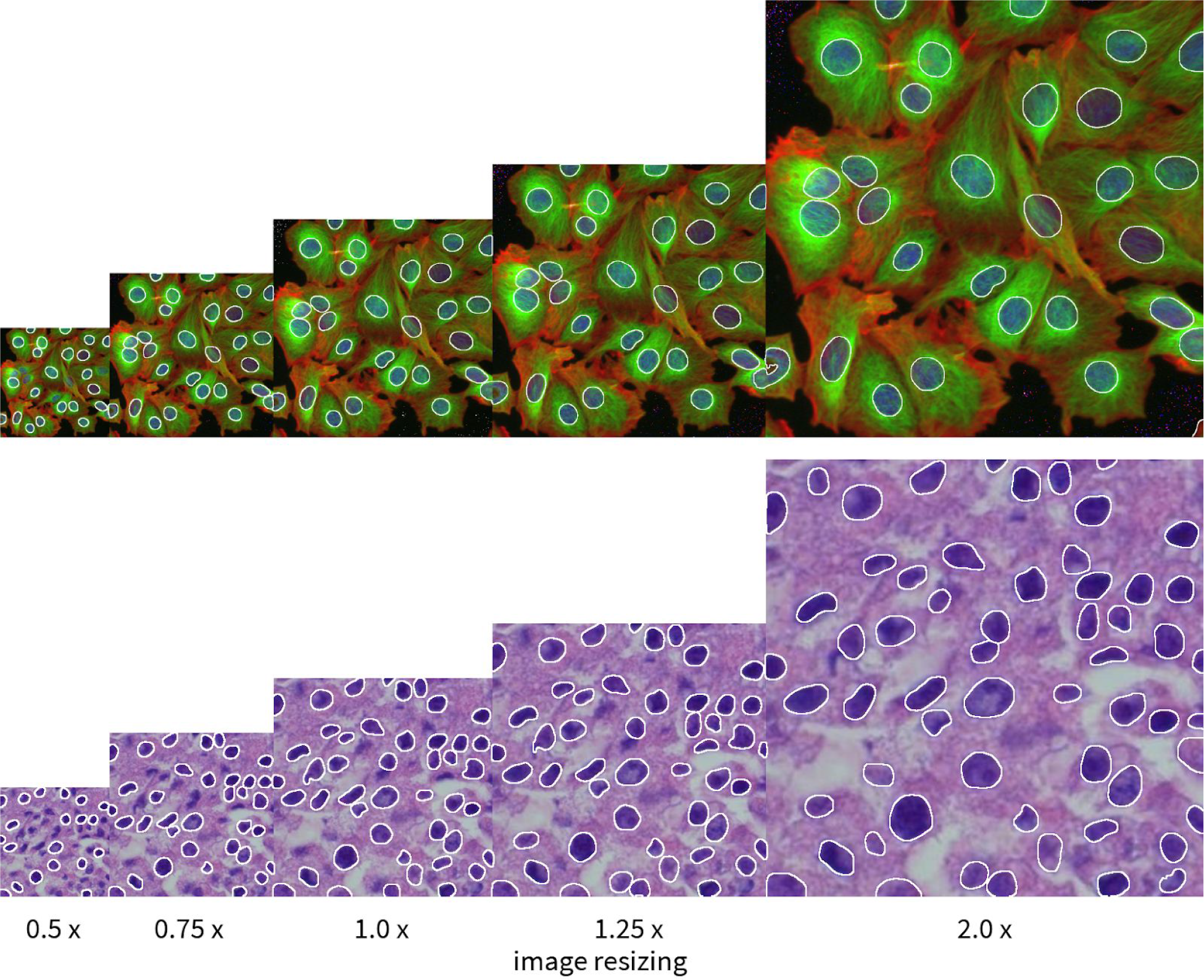
Mask R-CNN nucleus segmentations are reasonably robust to changes in scale, but we found that best performance can be achieved if nucleus size is fixed to 40 pixels diameter (at 1.0 scale factor).

#### 2.2. Inference

##### 2.2.1. Mask R-CNN prediction

The Mask R-CNN model trained as described above is used to predict segmentation masks when new images are provided as input. The images are resized before they are input to the network as described in Section 2.1.6.

##### 2.2.2. Post-processing and U-Net correction

We found that the segmentations could be further improved by postprocessing and refining nucleus contours using U-Net^13^. This encouraged better boundary reasoning between adjacent nuclei, and finer segmentations with the background. First, outlier objects were removed or merged as follows: 1) Smaller objects that were entirely within another object were eliminated. 2) objects that were surrounded by another object more than p_1_% were merged, and 3) objects smaller than p_2_ pixels area were removed. Next, U-Net based correction was performed (**Fig. S4**): 3) an optimal threshold p_3_ for U-Net probability values was determined, 4) a soft margin around the Mask R-CNN contour was defined for each object, with an extension of p_4_ pixels inwards and p_5_ outwards. The contour was extended/shrunk based on the U-Net predictions. 5) objects that had in total less than p6 mean U-Net probability were removed. Parameters p_1_.p_6_ were optimized on the training set. Best values were: (0.17, 44, 0.9375, 1, 1, 0.8).

**Figure S4.**
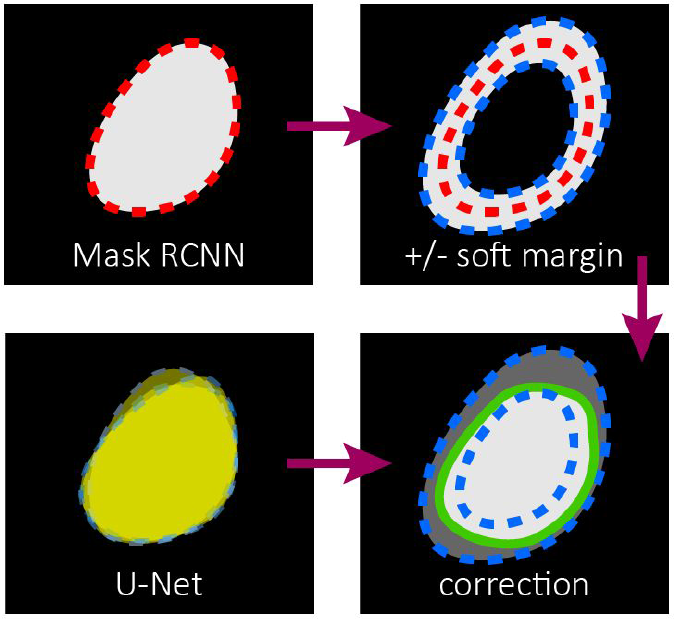
Post-processing contour correction with U-Net. Mask R-CNN segmentation contour was refined within a given maximum margin using U-Net segmentation. See Section 2.2.2 for details.

### 3. Kaggle competition

#### 3.1. Description

We designed our pipeline to recognize nuclei as accurately as possible in a wide variety of images acquired with different microscopes, under varying imaging conditions with different stains for nuclei of various cell types. This was the challenge set forth in the 2018 Data Science Bowl (DSB) by Kaggle, Booz Allen Hamilton and the Broad Institute. The competition included a preparatory stage 1, to which teams could submit their solutions during a four-month period and a 4-day long stage 2 final scoring period.

Existing nucleus segmentation methods do not generalize well, they perform well only on the limited experimental conditions they are designed or tuned for. The Data Science Bowl was highly successful in the sense that many robust solutions were developed that pushed the state-of-the-art in terms of segmentation performance and insensitivity to image type and quality. Solutions such as ours are now being developed into toolkits for biologists that will accelerate science by improving automation in identifying nuclei.

#### 3.2. Data

The official dataset for the challenge is composed of a training set and two tests sets, one for each stage. The number of images in each set is 670 (training), 65 (stage 1 test), and 3019 (stage 2 test), stage 1 test masks were released in the second stage. The final evaluation of the teams’ performance was measured on a subset of the stage 2 test set (the identity of the subset remained hidden to the competitors). Many of the competitors used additional data besides the provided training data, as this was permitted as long as participants shared their sources on the official competition website (https://www.kaggle.com/c/data-science-bowl-2018). Our annotated training data included 12 additional data sources besides the DSBdata, including some data sources annotated by experts in our institution. This extended the total number of training image/mask pairs from 735 to 1,102, and the number of annotated nuclei from 33,814 to 80,692 (not including the synthetic data). A summary of the data we used is provided in **Table S2**.

Using style transfer, we augmented our training data with synthetic image/mask pairs generated in the style of *k*=134 clusters of images from the DSB Stage 2 set, as described in Sections 2.1.2-2.1.4. This added 2,680 synthetic image/mask pairs to the training data (approximately 263,701 annotated nuclei).

We tested various versions of our method along with several competing methods on four test datasets: *DSB test1*, *DSB test2*, *fluo*, and *hist*. DSB test1 and DSB test2 are heterogeneous test sets from the Kaggle challenge (stage 1 and stage 2). The *fluo* dataset is fluorescence images of U2OS cells in a chemical screen taken from the Broad Bioimage Benchmark Collection (BBBC039)^9^. The *hist* dataset is a mixture of histology images collected from the internet and prostate H&E stained slides collected in-house.

A fraction of the histological images manually annotated in our lab were used as test set *hist* (see **Supplementary Data**). BBBC039 ^9^ images were used to train a fluorescent segmentation model, we refer as *fluo*. The *hist* and *fluo* test sets are disjoint from the respective training data.

#### 3.3. Evaluation metrics

The evaluation metric used for the DSB competition is based on the mean average precision^3^ at different intersection-over-union (IoU) thresholds. A successful nucleus detection was determined by an IoU test (also known as the Jaccard index):

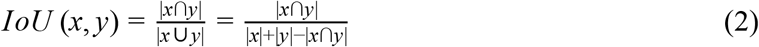

which measures the overlap between prediction pixels *x* and the annotation pixels *y* over the intersection of the two areas. Using a threshold ranging from 0.5 to 0.95, true positive (*TP*) detections, false positives (*FP*) detections and false negative (*FN*) detections were identified. For a threshold of 0.5, a predicted object is considered a “hit” if the IoU is greater than 0.5. For each threshold *t*, a a modified version of precision was calculated

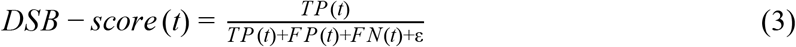

for all thresholds in (0.5, 0.95). These scores were averaged for all thresholds, and then the mean of the average scores is reported over the images in the test dataset. In addition to the DSB-score, we evaluated our results with three additional metrics based on the IoU detection test: mean average precision- (mAP), recall and F1-score. We used the same *t*, TP, FP and FN values as above. We also added a small ɛ = 10^−40^ value to the denominators.

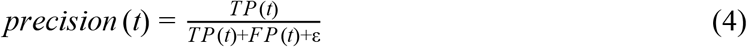

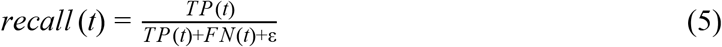

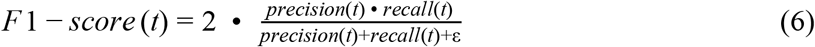

The same strategy was used to calculate mean values for these measures as was for the DSB-score, taking the average over various thresholds *t*, and the mean among the test images. In the following sections, we refer to these measures as mAP (mean average precision), mAR (mean average recall), and mF1 (mean average F1-score).

#### 3.4 Methods used for comparison

Our tests included several variations of our method along with six competing methods and several variations of our approaching using different style augmentation: *NOstyle* did not contain style augmented images, *AUTOstyle* used nuclei masks generated by the preseg network, and *GTstyle* used hand annotated ground truth to generate nuclei masks. CellProfiler (*CP*) ^1^ is a widely-used bioimage analysis software incorporating several methods to segment, measure and analyze cellular compartments. We created multiple pipelines for the different image types of the test sets – except for our fluorescent set which comprised of a single experiment. *Preseg* refers to our general scale-independent pre-segmentation model while *postComp* is our final refined post-competition submission (an AUTOstyle model customized for DSB test2).

We compared against several other approaches including *ilastik ^14^*, which provides a pixel classification setup where users can manually annotate regions of the input images to desired classes and obtain predictions as either probability maps or segmented images. Segmentations were obtained by applying a threshold to probabilities from *ilastik* (with additional object splitting). *Unet4nuclei* ^9^ is an implementation of the popular U-Net deep learning approach to segmentation. *GVF* ^15^, or gradient vector flow, is an active contour-based segmentation method suitable if objects are bright regions on a dark background. Pipelines of these compared methods are provided in **Supplementary Software 2**. *DSB1* and *DSB2* are the first and second place entries on the final Kaggle leaderboard. The approach from *DSB1* (https://www.kaggle.com/c/data-science-bowl-2018/discussion/54741) uses a very deep U-Net architecture along with prediction of touching borders. *DSB2* also uses a U-Net approach, and forces the network to predict relative locations within each nucleus (https://github.com/jacobkie/2018DSB).

#### 3.5. Results

We tested the methods outlined in Section 3.4 on four test datasets: *DSB test1*, *DSB test2*, *fluo*, and *hist*, described in Section 3.2. The resulting DSB-scores are presented in Table S1. When running these tests, the test data was never included in the data to train the model, e.g. when testing on *DSB test1*, the *DSB test1* data was held out from the training set. Similarly, when testing on hist, *biomag2* and *biomag6* subsets were held out.

**Table S1.**
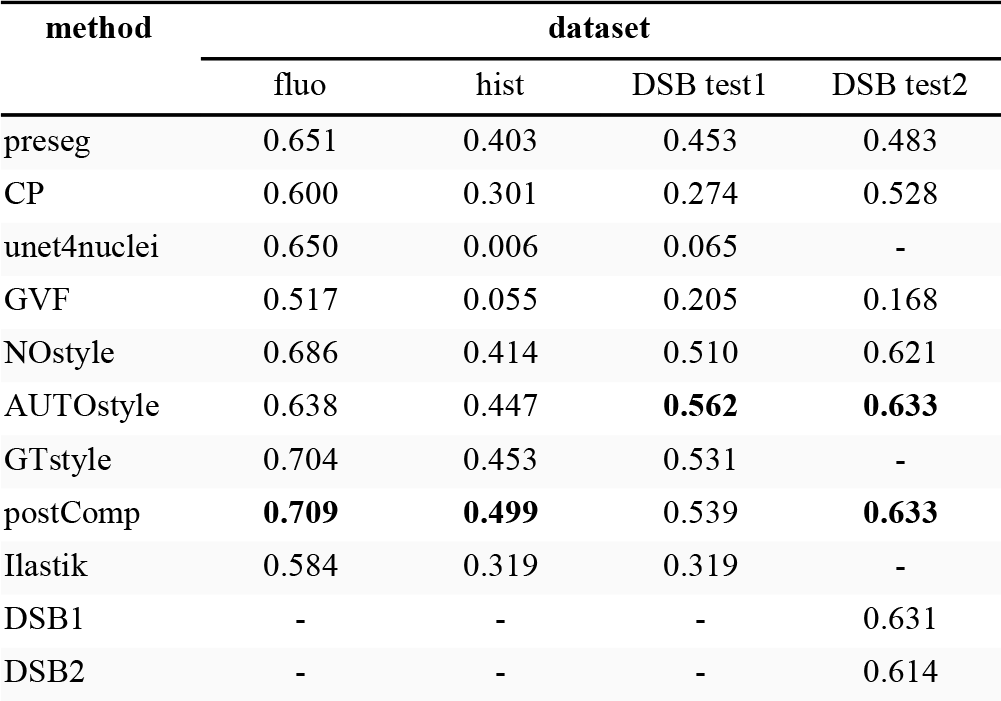
DSB scores. We tested the methods described in Section 3.4 on four test datasets: *DSB test1*, *DSB test2*, *fluo*, and *hist. Fluo* was the BBBC039 ^9^ image set of 200 DAPI-labelled fluorescent images while *hist* was a randomly selected set of 50 in-house labeled histological images. DBS test1 and test2 are the official test sets of the DSB 2018 competition. The DSB-score is the mean average precision at different intersection-over-union (IoU) thresholds (see Section 3.3). *GTStyle* is not reported for DSB test2 because annotations are not available, as a result the *postComp* model is the same as *AUTOstyle*. The best performing method’s score is highlighted in bold for each test set.

**Figure S5.**
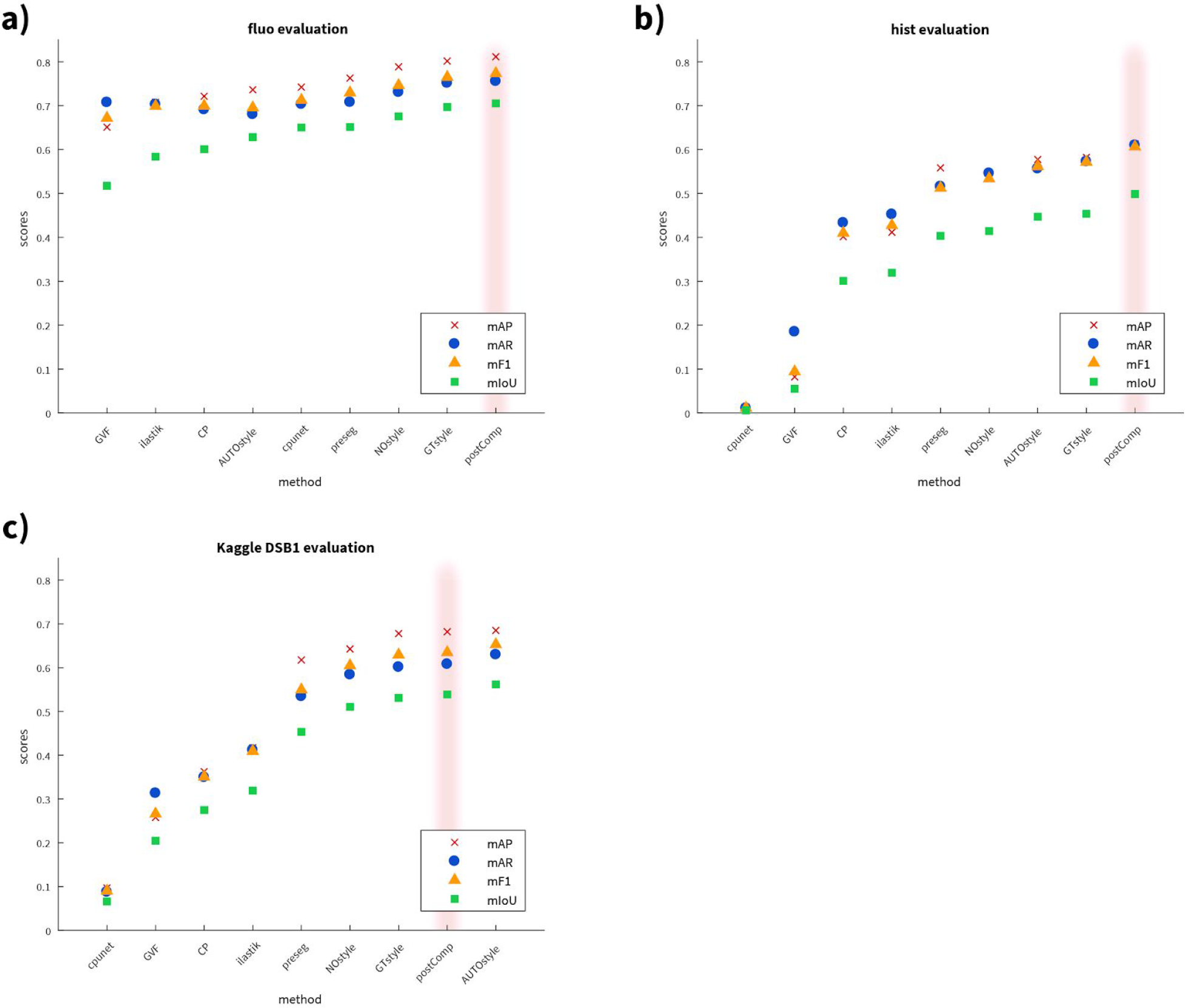
Evaluation of the methods on *fluo*, *hist*, and *DSB test1* using different metrics: mAP: mean average precision, mAR: mean average recall, mF1: mean average F1-score, mIoU: mean intersection over union. See definitions in 3.3. a) test set *fluo*, b) test set *hist*, c) test set *DSB test1*. *postComp* is highlighted in light pink shade. Methods are sorted by mIoU. Results on DSB test2 are not reported because ground truths are not available.

The results demonstrate that our style transfer approach improves performance in test sets containing data from heterogeneous sources: *hist*, *DSB test1* and *DSB test2*. We also see excellent performance on single domain fluorescence data, *fluo*. Comparing the results of our method with and without style transfer augmentation, we see a clear trend towards increased performance with style transfer augmentation. If we have access to ground truth nucleus masks (*GTstyle*) our performance improves, though in many realistic scenarios such masks will not be available. **Figure S2** shows the output of the various methods we tested on challenging examples (note that *DSB1* and *DSB2* are not reported because we did not have access to their code). In **Figure S5**, we present mAP, mAR, mF1 and mIoU metrics for the various methods on each dataset. As expected, there is a strong correlation between the metrics.

### 4. Computational environment

#### 4.1. Software

Our pipeline is implemented using a shell script to allow continuous execution of the entire pipeline. Python 3 scripts execute the training and inference of Mask R-CNN, U-Net, and pix2pix which rely on the TensorFlow, Keras, and PyTorch environments. The clustering, post-processing, and initial steps of style transfer are implemented in Matlab. Our software is available for download at: https://github.com/spreka/biomagdsb.

The entire pipeline can be run both under Linux and Windows. In a typical use case, it is not necessary to retrain any of the models. Calling the *postComp* method without post processing provides excellent results. Alternatively, an online version of our method is available at www.nucleAIzer.org.

#### 4.2. Hardware

Our methods were trained and tested on a variety of Nvidia graphics cards, including GTX 1070, 1080Ti, and Titan Xp.

### 5. Related work

#### 5.1. Mask R-CNN

Kaming He et al. ^12^ published Mask R-CNN as an extension of Faster R-CNN to allow simultaneous instance detection and segmentation. The network architecture is similar to that of Faster R-CNN: feature extraction uses ResNet (50 or 101 layers) or alternatively Feature Pyramid Network (FPN), while head is as in Faster R-CNN extended with a fully convolutional mask prediction branch. A detailed discussion of extended R-CNN versions can be found in ^20^.

#### 5.2. U-Net

U-Net ^13^ was specifically created for bioimage segmentation with an encoder-decoder architecture and skip connections between layers of the encoding branch and decoding branch to provide the decoder with access to spatial information to reason about upsampling the segmentation.

## Additional supplementary info

**Supplementary Table 2.:** https://docs.google.com/document/d/1DhtWss5zFskrDLU9Isqs556ygBRtxgRkZdUJlJX_OtI/edit?usp=sharing

**Supplementary Data 1.:** https://drive.google.com/file/d/1x_-EM8q4Mawu-GEV02Y6kFTuw8EznJGU/view?usp=sharing

**Supplementary Software 1.:** https://github.com/spreka/biomagdsb.

**Supplementary Software 2:** https://drive.google.com/file/d/111aWsov8nuyLSGx1TDnT7XyV8-BFRZRw/view?usp=sharing

## Footnotes

1 Our method achieved the top-score after the competition ended. An early version of our approach placed 27th out of 739 submissions in round 2 of the competition.

2 As defined on the competition website.

3 As defined on the competition website

