## Supplementary figures and tables for "A deep learning framework for nucleus segmentation using image style transfer"

### Supplementary figures

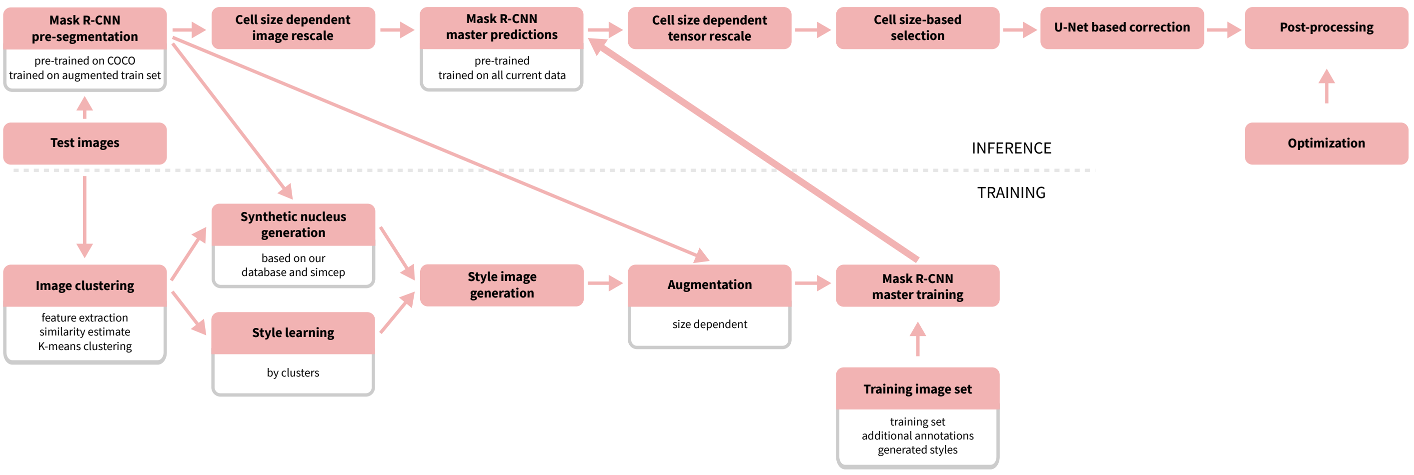


**Figure S1.** A detailed overview of the proposed pipeline.

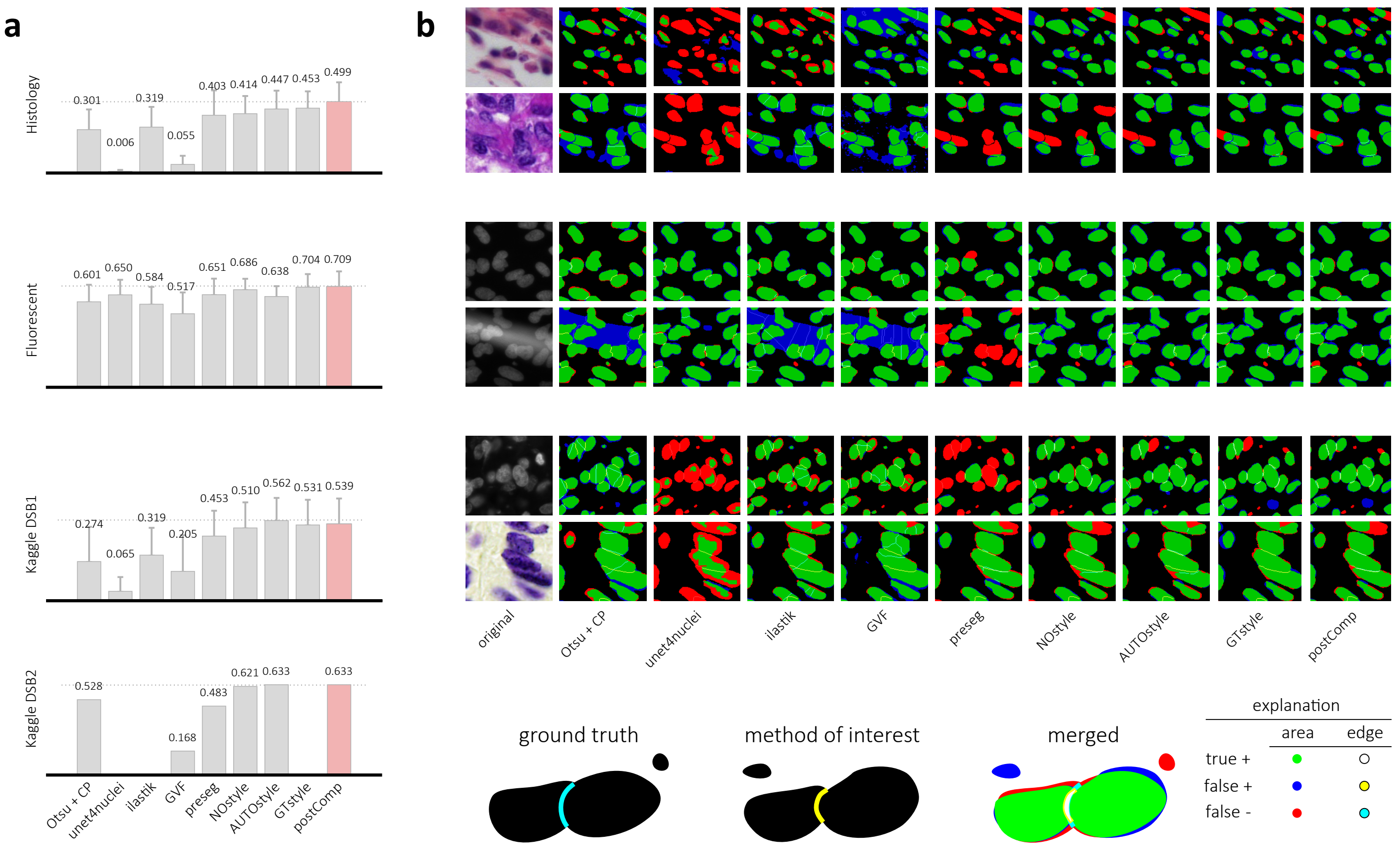


**Figure S2.** Results from Figure 2 in more detail. a) DSB scores with error bars for the image sets tested. b) Segmentation results for various methods. preseg, NOstyle, AUTOstyle, GTstyle and postComp refer to variations of our approach. See sections 3.2. and 3.4-3.5. for details.

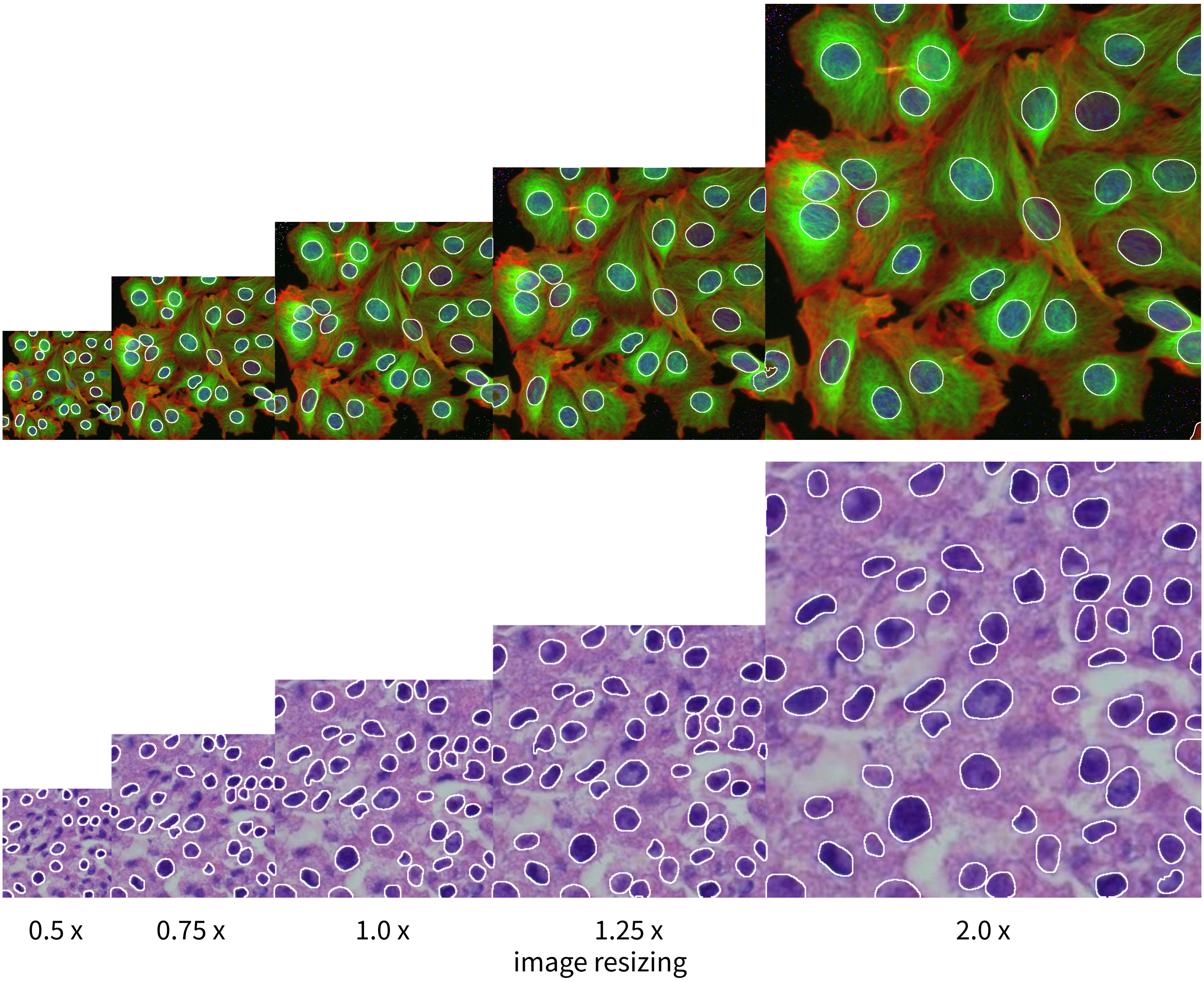


**Figure S3.** Mask R-CNN nucleus segmentations are reasonably robust to changes in scale, but we found that best performance can be achieved if nucleus size is fixed to 40 pixels diameter (at 1.0x magnification).

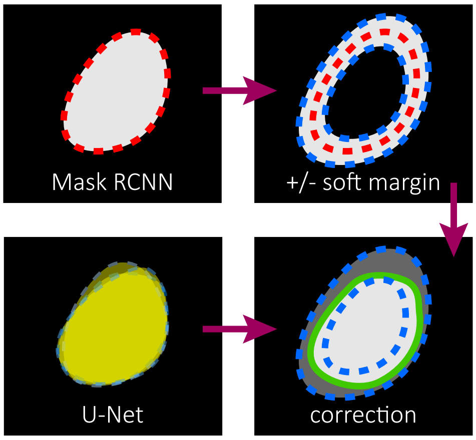


**Figure S4.** Post-processing contour correction with U-Net. Mask RCNN segmentation contour was refined within  a given maximum margin using U-Net segmentation. See Section 2.2.2 for details.

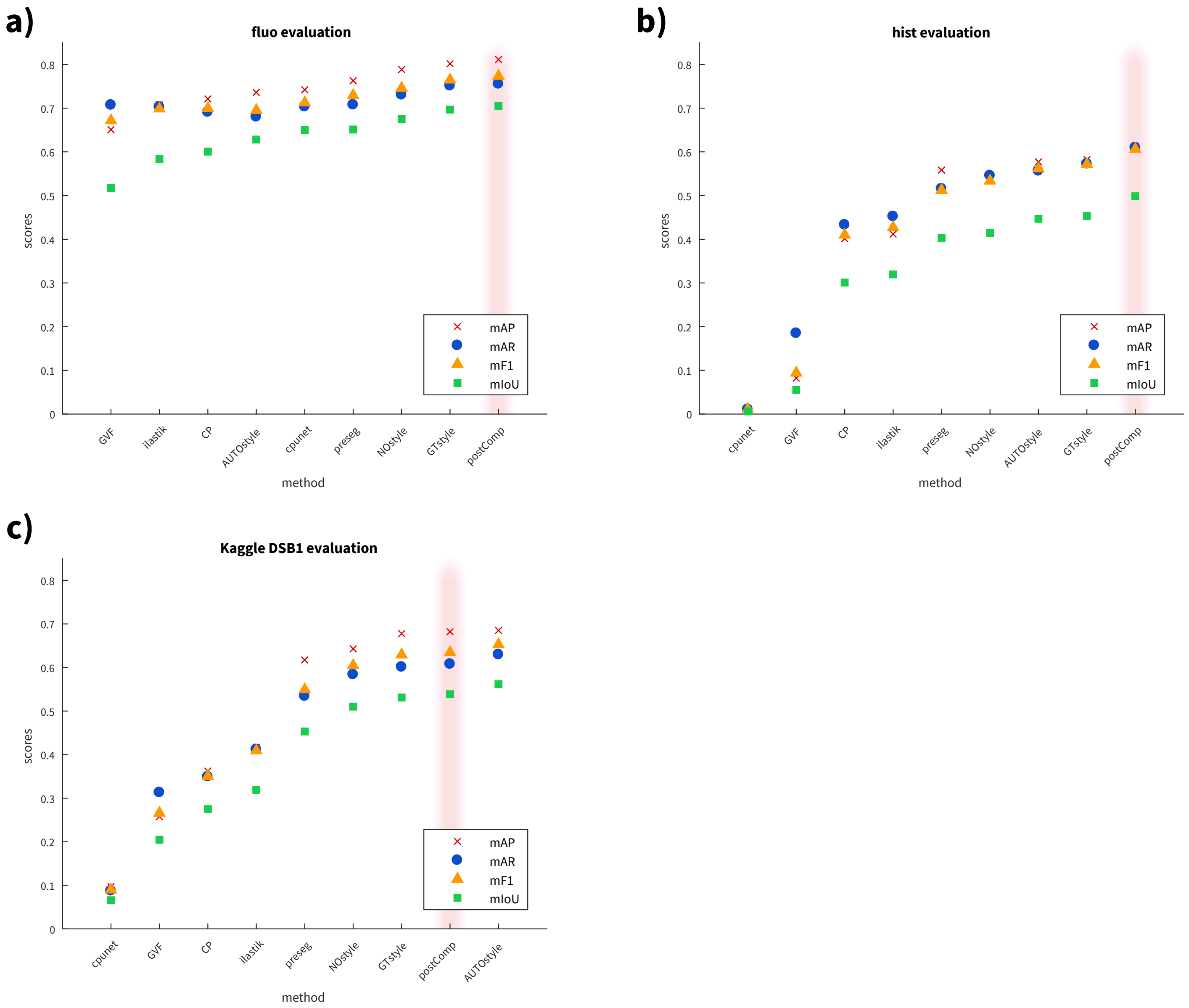


**Figure S5.** Evaluation of the methods on fluo, hist, and DSB test1 using different metrics: mAP: mean average precision, mAR: mean average recall, mF1: mean average F1-score, mIoU: mean intersection over union. See definitions in 3.3. a) test set fluo, b) test set hist, c) test set DSB test1. postComp is highlighted in light pink shade.

### Supplementary tables

**Table S1: DSB scores**

| **method** | **dataset** | | | |
|---|---|---|---|---|
| | fluo | hist | DSB test1 | DSB test2 |
| preseg | 0.651 | 0.403 | 0.453 | 0.483 |
| CP | 0.600 | 0.301 | 0.274 | 0.528 |
| unet4nuclei | 0.650 | 0.006 | 0.065 | - |
| GVF | 0.517 | 0.055 | 0.205 | 0.168 |
| NOstyle | 0.686 | 0.414 | 0.510 | 0.621 |
| AUTOstyle | 0.638 | 0.447 | **0.562** | **0.633** |
| GTstyle | 0.704 | 0.453 | 0.531 | - |
| postComp | **0.709** | **0.499** | 0.539 | **0.633** |
| Ilastik | 0.584 | 0.319 | 0.319 | - |
| Kaggle1 | - | - | - | 0.631 |
| Kaggle2 | - | - | - | 0.614 |

**Table S1.** DSB scores. We tested the methods described in Section 3.4 on four test datasets: DSB test1, DSB test2, fluo, and hist. Fluo was the BBBC039 9 image set of 200 DAPI-labelled fluorescent images while hist was a randomly selected set of 50 in-house labeled histological images. DBS test1 and test2 are the official test sets of the DSB 2018 competition. The DSB-score is the mean average precision at different intersection-over-union (IoU) thresholds (see Section 3.3). The DSB test2 set has both AUTOstyle and postComp models with the highest score because in this specific case the automated style transfer learning was derived from test2, that was our final competition model. The best performing method's score is highlighted in bold for each test set.

**Table S2: Training dataset statistics**

| property | | subset | | | | | | | | | | | | | |
|---|---|---|---|---|---|---|---|---|---|---|---|---|---|---|---|
| | | DSB train | DSB test1 | Weebly | TNBC | BBBC021 | Internet | biomag 02 | biomag 04 | biomag 05 | biomag 06 | jw | ISBI 2009 | AS | 00733 |
| image type | fluo normal | x | x | - | - | x | x | - | - | - | - | x | x | x | - |
| | fluo clumped | x | x | - | - | x | x | - | x | x | x | x | - | x | x |
| | tissue simple | x | x | - | - | - | x | x | - | - | - | - | - | - | - |
| | tissue complex | x | x | x | x | - | x | - | - | - | x | - | - | - | - |
| image size | min | 256x256 | 256 256 | 1000x1000 | 512x512 | 1280x1024 | 213x65 | 812x828 | 288x286 | 256x256 | 256x340 | 956x1242 | 1030x1349 | 512x512 | 512x512 |
| | max | 1040x1388 | 520 696 | - | - | - | 2848x4272 | 1024x1344 | 326x327 | 1024x1360 | 2048x2048 | 1040x1388 | - | - | - |
| number of images | | 670 | 65 | 30 | 50 | 5 | 97 | 11 | 4 | 5 | 32 | 20 | 48 | 11 | 54 |
| number of nuclei | mean/image | 44.27 | 4152 | 565.23 | 81.12 | 124.40 | 99.18 | 94.27 | 53.25 | 197.60 | 67.03 | 42.30 | 38.10 | 110.27 | 136.15 |
| | total | 29662 | 63.88 | 16957 | 4056 | 622 | 9620 | 1037 | 213 | 988 | 2145 | 846 | 1829 | 1213 | 7352 |
| models trained on | | all | all but this | all | all | all | all but hist | all but hist | all | all | all but hist | all | all | all | all |
| models tested on | | - | DSB test1 | - | - | - | hist | hist | - | - | hist | - | - | - | - |
| data source | | DSB | DSB | 1 | 2 | 3, 4 | 5, 6 | 7 | 8 | 9 | 10 | 11 | 11, 12 | 11 | 11 |

**Table S2.** Statistics about the training data. Data sources in the last row are as follows.
1: https://nucleisegmentationbenchmark.weebly.com/dataset.html
2: Naylor Peter Jack, Walter Thomas, Laé Marick, & Reyal Fabien. (2018). Nuclei Segmentation in Histopathology Images Using Deep Neural Networks (Version 1.0) [Data set]. Zenodo. http://doi.org/10.5281/zenodo.1174353
3: http://mct.aacrjournals.org/content/9/6/1913
4: https://www.nature.com/articles/nmeth.2083
5: https://www.google.hu/search?q=histology+microscope+nucleus&-source=lnms&tbm=isch&sa=X&ved=0ahUKEwiIxb6v_J3aAhXFFJoKHYdzAxUQ_AUICigB&biw=1366&bih=654
6: https://www.google.hu/search?q=dapi+stained+nuclei&client=-firefox-b-ab&dcr=0&source=lnms&tbm=isch&sa=X&ved=0ahUKEwik4bK8_Z3aAhXLDZoKHZXTDZkQ_AUICigB&biw=1366&bih=654
7: https://www.nature.com/articles/nmeth.3323
8: https://www.ncbi.nlm.nih.gov/pubmed/28122742
9: https://www.nature.com/articles/s41467-017-02628-4
10: Peter Horvath Laboratory
11: shared by other participating teams of the DSB on the competition website
12: https://www.ncbi.nlm.nih.gov/pmc/articles/PMC2901896/
